# C-di-AMP accumulation disrupts glutathione metabolism and inhibits virulence program expression in *Listeria monocytogenes*

**DOI:** 10.1101/2024.01.18.576247

**Authors:** Cheta Siletti, Matthew Freeman, Zepeng Tu, David M. Stevenson, Daniel Amador-Noguez, John-Demian Sauer, TuAnh N. Huynh

## Abstract

C-di-AMP is an essential second messenger in many bacteria but its levels must be regulated. Unregulated c-di-AMP accumulation attenuates the virulence of many bacterial pathogens, including those that do not require c-di-AMP for growth. However, the mechanisms by which c-di-AMP regulates bacterial pathogenesis remain poorly understood. In *Listeria monocytogenes*, a mutant lacking both c-di-AMP phosphodiesterases, denoted as the ΔPDE mutant, accumulates a high c-di-AMP level and is significantly attenuated in the mouse model of systemic infection. All key *L. monocytogenes* virulence genes are transcriptionally upregulated by the master transcription factor PrfA, which is activated by reduced glutathione (GSH) during infection. Our transcriptomic analysis revealed that the ΔPDE mutant is significantly impaired for the expression of virulence genes within the PrfA core regulon. Subsequent quantitative gene expression analyses validated this phenotype both at the basal level and upon PrfA activation by GSH. A constitutively active PrfA^*^ variant, PrfA G145S, which mimics the GSH-bound conformation, restores virulence gene expression in ΔPDE but only partially rescues virulence defect. Through GSH quantification and uptake assays, we found that the ΔPDE strain is significantly depleted for GSH, and that c-di-AMP inhibits GSH uptake. Constitutive expression of *gshF* (encoding a GSH synthetase) does not restore GSH levels in the ΔPDE strain, suggesting that c-di-AMP inhibits GSH synthesis activity or promotes GSH catabolism. Taken together, our data reveals GSH metabolism as another pathway that is regulated by c-di-AMP. C-di-AMP accumulation depletes cytoplasmic GSH levels within *L. monocytogenes* that leads to impaired virulence program expression.

**IMPORTANCE:** C-di-AMP regulates both bacterial pathogenesis and interactions with the host. Although c-di-AMP is essential in many bacteria, its accumulation also attenuates the virulence of many bacterial pathogens. Therefore, disrupting c-di-AMP homeostasis is a promising antibacterial treatment strategy, and has inspired several studies that screened for chemical inhibitors of c-di-AMP phosphodiesterases. However, the mechanisms by which c-di-AMP accumulation diminishes bacterial pathogenesis are poorly understood. Such understanding will reveal the molecular function of c-di-AMP, and inform therapeutic development strategies. Here, we identify GSH metabolism as a pathway regulated by c-di-AMP that is pertinent to *L. monocytogenes* replication in the host. Given the role of GSH as a virulence signal, nutrient, and antioxidant, GSH depletion impairs virulence program expression and likely diminishes host adaptation.

## INTRODUCTION

The Gram-positive bacterial pathogen *Listeria monocytogenes* is a leading cause of mortality and hospitalization among foodborne illnesses. Although acquired orally, *L. monocytogenes* can cross the intestinal barrier to cause systemic infection with mortality rates approaching 16% in clinical cases (1). A hallmark of *L. monocytogenes* pathogenesis is its intracellular life cycle (2). *L. monocytogenes* can invade many mammalian cell types, including non-phagocytic cells, using surface-anchored internalins such as InlA and InlB. Following host-cell entry, *L. monocytogenes* escapes the vacuole using the pore-forming toxin listeriolysin O (LLO, encoded by the *hly* gene), the phospholipases PlcA and PlcB, and the metalloprotease Mpl. In the cell cytosol, *L. monocytogenes* uses ActA to polymerize host actin, required for intracellular motility and cell-to-cell spread. All virulence genes required for the intracellular lifestyle, including those listed above, are transcriptionally regulated by the master transcription factor PrfA (3).

PrfA belongs to the Crp/Fnr family of transcription factors, which function as homodimers with a DNA-binding helix-turn-helix motif in each monomer (3, 4). Unlike other Crp/Fnr proteins that require a co-factor for DNA binding, apo-PrfA can bind to its consensus palindromic DNA operator, called PrfA box, to maintain a basal expression of some virulence genes in the absence of activating signals. During infection, PrfA is allosterically activated by reduced glutathione (GSH), which is derived from the host or synthesized by *L. monocytogenes* (5). GSH binds PrfA at the stoichiometry of one GSH per PrfA monomer, inducing an active DNA-binding conformation of the helix-turn-helix motif (6). A single mutation (G145S) near the GSH binding site renders PrfA constitutively active by mimicking the GSH-bound conformation (6, 7). During broth growth in *Listeria* Synthetic Medium (LSM), PrfA can be activated by supplementation with GSH or reducing reagents such as TCEP (8).

*L. monocytogenes* makes and secretes c-di-AMP, a nucleotide second messenger that regulates many molecular targets in bacterial cells (9, 10). C-di-AMP-binding proteins within *L. monocytogenes* have different cellular functions, such as potassium and carnitine uptake, central metabolism and ppGpp synthesis, and signal transduction (11–15). During infection, secreted c-di-AMP is recognized by mammalian cytosolic receptors to activate inflammatory and type I interferon responses (9, 16). Type I interferon response is considered to promote *L. monocytogenes* pathogenesis during systemic infection, but activates an anti-bacterial response in the gastrointestinal tract (17, 18).

C-di-AMP homeostasis is critical to *L. monocytogenes* growth and infection. In *L. monocytogenes*, c-di-AMP is synthesized from ATP by a single diadenylate cyclase, DacA, and degraded by the phosphodiesterases PdeA and PgpH into the linear nucleotide pApA (19). In rich media, such as Brain Heart Infusion (BHI) broth, the Δ*dacA* mutant accumulates a toxic level of ppGpp that inhibits bacterial growth (20). By contrast, the Δ*pdeA* Δ*pgpH* mutant (hereafter denoted as the ΔPDE mutant) does not exhibit an appreciable growth defect in BHI, but is greatly attenuated for virulence in the mouse model of intravenous infection, despite hyper-activating type I interferon response (21). These phenotypes indicate that the mechanisms underlying ΔPDE virulence attenuation are related to bacterial defects during infection.

The pathways and molecular targets leading to diminished pathogenesis at high c-di-AMP levels are not clearly defined. Here, we found that c-di-AMP accumulation inhibits the expression of the PrfA core regulon in *L. monocytogenes*, and this defect contributes to virulence attenuation in the ΔPDE strain. We further found that c-di-AMP accumulation impairs GSH uptake, and either inhibits GSH synthesis or promotes GSH catabolism. Combined, these defects result in a cytoplasmic GSH depletion that impairs PrfA function. Our findings reveal GSH metabolism as another pathway that is regulated by c-di-AMP, with implications in *L. monocytogenes* virulence program and nutrient acquisition inside the host.

## RESULTS

### The ΔPDE mutant is defective for virulence gene expression in the PrfA core regulon

The ΔPDE mutant was previously shown to be highly attenuated for virulence in a mouse model of infection (21). To identify genes that could contribute to ΔPDE virulence defect, we performed RNAseq to profile the transcriptomes of the wild-type (WT) and ΔPDE strains, grown in LSM to mid-exponential phase (20). The cDNA libraries were sequenced at the depth of 17 – 20 million reads, and had 856x-1042x mapped read coverage of the *L. monocytogenes* genome. Compared to WT, the ΔPDE strain displayed mostly a transcriptional downshift, with 67 genes being down-regulated by ≥4-fold, and 9 genes being up-regulated by ≥4-fold (P value < 0.05) (**Table S1**). Remarkably, the vast majority of down-regulated genes in ΔPDE belong to the 10403S prophage locus (41 genes) or the PrfA core regulon (11 genes) (**Fig. 1A**). In addition to the core regulon, PrfA has been shown to putatively or indirectly regulate the expression of up to 145 other genes (22). We did not observe those genes among the most significantly altered in ΔPDE, except for the upregulation of a cellobiose transporter (*lmo2683*) and two genes of hypothetical function (*lmo0186* and *lmo0019*) **(Table S1)**. Furthermore, *kdpA* expression was also reduced by ∼6-fold in the ΔPDE strain, consistent with an inhibitory effect of c-di-AMP on the expression of the *kdp* operon, which encodes a potassium uptake system (**Table S1**) (13, 23).

**Figure 1:**
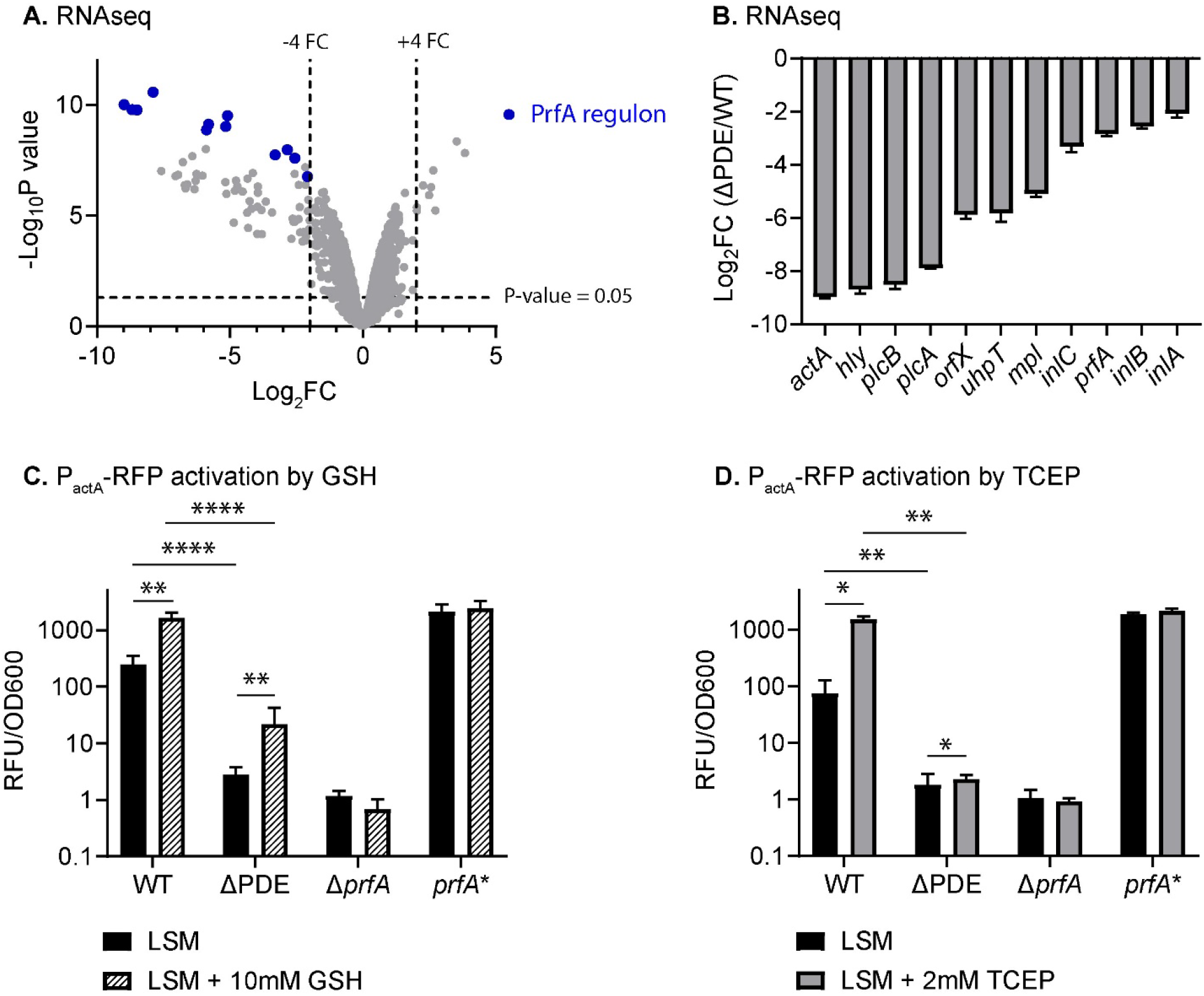
C-di-AMP accumulation impairs virulence gene expression in *L. monocytogenes*. **A**. Volcano plot of differentially expressed genes in the ΔPDE mutant compared to the WT strain. RNAseq was performed on mid-log cultures grown in *Listeria* Synthetic Medium (LSM). Blue dots indicate genes within the PrfA core regulon. **B**. Log_2_fold change of PrfA-regulated gene expression was calculated from read counts per million (cpm) of two independent ΔPDE cultures, compared to the average of two independent WT cultures. Data in A and B are from the same RNAseq experiment. **C**. PrfA-regulated gene expression was quantified in *L. monocytogenes* strains carrying P_actA_::RFP transcriptional reporter. The Δ*prfA* and *prfA*^*^ strains were examined as negative and positive controls, respectively. Cultures were grown to mid-log in LSM or LSM + 10mM GSH. Red fluorescence was normalized to OD600 of each respective culture. **D**. Red fluorescence was quantified as in C. LSM cultures were left untreated or treated with 2mM TCEP for 2 hours. Data in C and D are average of 3-6 independent experiments. Error bars show standard deviations. Statistical analyses were performed by two-way ANOVA with multiple comparisons for the indicated pairs: ns, non-significant; ^*^, P < 0.05; ^**^, P < 0.01; ^****^, P < 0.0001

PrfA is the master transcription factor that upregulates *L. monocytogenes* virulence gene expression required for the intracellular life cycle (4). Our RNAseq analysis revealed that the PrfA core regulon was significantly down-regulated in the ΔPDE strain (**Fig. 1A-B**). To validate RNAseq results, we employed RT-qPCR to quantify the expression of three genes in the PrfA core regulon: *prfA, hly* (an early gene), and *actA* (a late gene). In LSM cultures, the ΔPDE strain was significantly impaired for the expression of all three genes, reflecting a defect in basal PrfA function (**Fig. S1A-C**). To examine gene expression upon PrfA activation, we next performed RT-qPCR for cultures grown in the presence of GSH. Although these genes were up-regulated by GSH in both the WT and ΔPDE strains, their expression levels in ΔPDE remained significantly lower than in WT (**Fig. S1A-C**). Furthermore, of the three genes, we noticed that the ΔPDE strain was most impaired for *actA* expression, which was reduced by 10 – 100-fold compared to the basal and activated levels in WT (**Fig. S1C**). As an independent method to quantify *actA* expression, we used a red fluorescent reporter fused with the *actA* promoter (P_actA_-RFP) (8). This assay confirmed that the ΔPDE strain was significantly defective for *actA* expression both at the basal level and upon PrfA activation by GSH or TCEP (**Fig. 1C-D**).

### Defective PrfA function contributes to virulence attenuation at high c-di-AMP levels

To assess whether a defect in PrfA function contributes to ΔPDE virulence attenuation, we replaced the native *prfA* allele with a constitutive PrfA^*^ variant (G145S) that structurally mimics the GSH-bound form (6, 7). In transcriptional reporter assays with P_actA_-RFP, the *prfA*^*^ strain exhibited constitutive *actA* expression, independent of GSH supplementation, as expected (**Fig. 2A**). In the ΔPDE background, the *prfA*^*^ allele restored *actA* expression to the WT level in LSM + GSH (**Fig. 2A**). Interestingly, *actA* expression was not constitutive in ΔPDE *prfA*^*\**^. These data indicate that impaired PrfA activity is responsible for low virulence gene expression in ΔPDE. Furthermore, there appears to be factors that inhibit PrfA activity in the absence of GSH.

**Figure 2:**
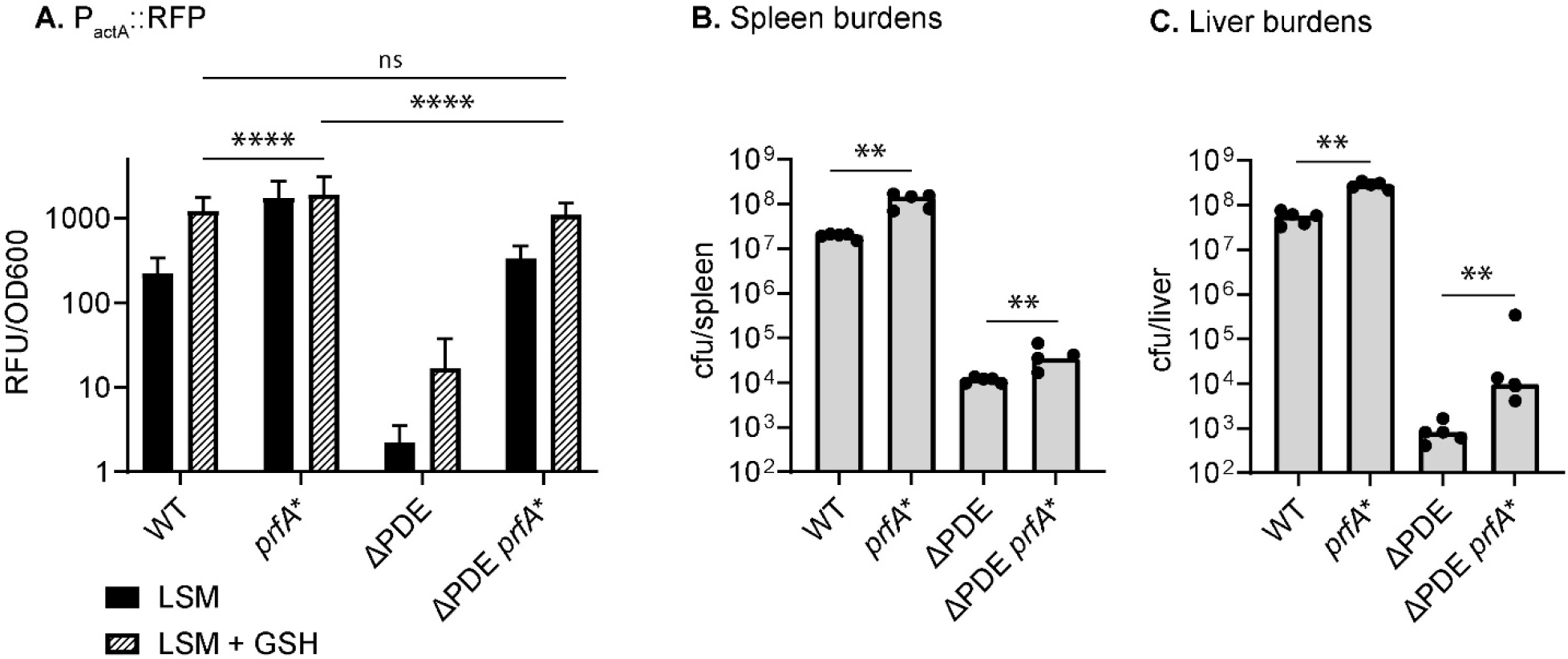
A constitutive PrfA variant (PrfA^*^) restores virulence gene expression and partially rescues virulence defect upon c-di-AMP accumulation. **A**. P_actA_::RFP transcriptional activity was quantified in WT, ΔPDE, and isogenic strains in which the native *prfA* allele is replaced with *prfA*^*^ (G145S). Red fluorescence, normalized to OD600, was measured for mid-log cultures grown in LSM or LSM + 10mM GSH, and normalized to OD600. Error bars show standard deviations. **B-C**. Bacterial burdens in the spleen and liver at 48 hours post-intravenous infection with 1 x 10^5^ cfu of *L. monocytogenes*. Each dot represents *L. monocytogenes* burden from one animal. Bars represent medians of the groups. Statistical analyses were performed by two-way ANOVA in A and Mann-Whitney test in B, with multiple comparisons for the indicated pairs: ns, non-significant; ^**^, P < 0.01; ^****^, P < 0.0001.

We next examined the contribution of PrfA defect to ΔPDE virulence attenuation in a mouse model of intravenous infection. In the WT background, *prfA*^*^ increased bacterial burdens in both the spleen and liver, as expected (**Fig. 2B-C**). Compared to ΔPDE burdens, the median burdens of ΔPDE *prfA*^*^ were increased by ∼10-fold in the liver, and by ∼3-fold in the spleen (**Fig. 2B-C**). Thus, a defect in PrfA function partially contributes to ΔPDE virulence attenuation in the liver. Additionally, our data also reveal that there are additional mechanisms underlying the virulence defect of ΔPDE at high c-di-AMP levels.

### Glutathione deficiency impairs PrfA activity in the ΔPDE strain

The partial restorative effect of PrfA^*^ in virulence suggests that the ΔPDE strain might be impaired for PrfA activation by GSH. We found that GSH levels were significantly depleted in the ΔPDE strain compared to WT, in all culture conditions that we quantified PrfA-regulated gene expression (LSM, LSM + GSH, and LSM + TCEP) (**Fig. 3A-B)**. The deficiency in GSH was not due to oxidation, since the total glutathione pool and oxidized glutathione (GSSG) were also reduced in ΔPDE (**Fig. S2A-B)**. Furthermore, a previous study found that methylglyoxalase is important to maintain a sufficient GSH pool in *L. monocytogenes* (24). We found no evidence for an impaired methylglyoxalase activity in the ΔPDE strain, since ΔPDE exhibited a comparable methylglyoxal susceptibility to WT (**Fig. S2C**).

**Figure 3:**
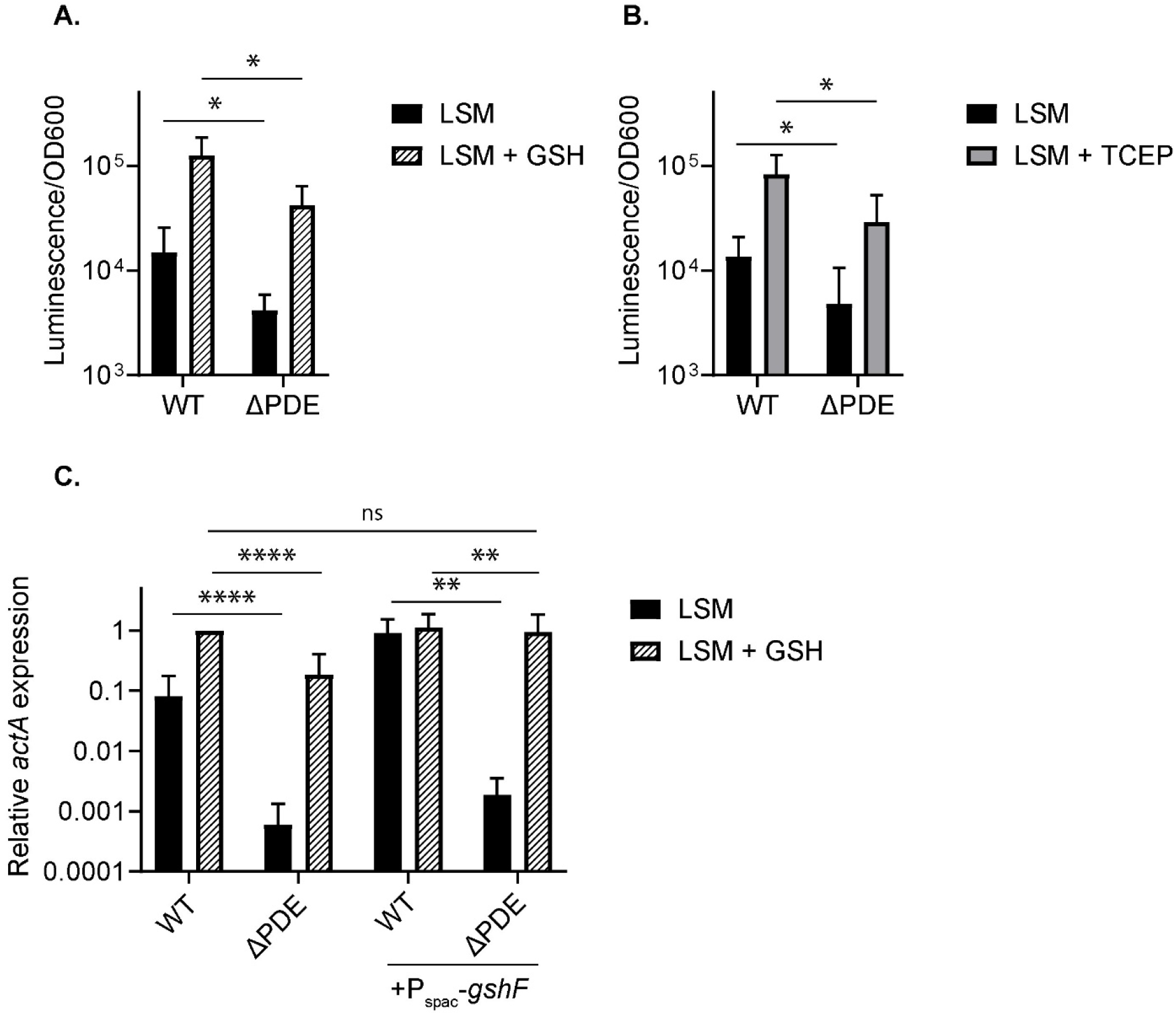
A deficiency in reduced glutathione (GSH) contributes to diminished virulence gene expression at high c-di-AMP levels. **A**. Reduced glutathione (GSH) levels of mid-log cultures grown in LSM or LSM + 10mM GSH. GSH levels are reported by luminescence and normalized to OD600. Several GSH standards were quantified to determine the linear range of the assay, and all biological samples were quantified at appropriate dilutions to ensure measurements were within the linear range. **B**. Mid-log LSM cultures were left untreated, or treated with 2mM TCEP for 2 hours, and quantified for GSH levels as in A. **C**. *actA* gene expression by RT-qPCR. Cultures were grown in LSM or LSM + 10mM GSH to mid-log phase. In each culture, the expression level of *actA* was normalized to that of *rplD* as a housekeeping gene. Error bars represent standard deviations. Statistical analyses were performed by two-way ANOVA with multiple comparisons for the indicated pairs: ns, non-significant; ^*^, P < 0.05; ^**^, P < 0.01; ^****^, P < 0.0001

*L. monocytogenes* synthesizes GSH by the glutathione synthase GshF (25), and also imports GSH via CtaP and OppDF (26). The ΔPDE strain could fully induce *actA* expression to the activated level in WT, but only upon both GSH supplementation and *gshF* over-expression (**Fig. 3C**). These data suggest that GSH deficiency impairs PrfA function in the ΔPDE strain.

### C-di-AMP accumulation inhibits GSH uptake

We sought to determine how GSH is depleted in the ΔPDE mutant. We evaluated GSH uptake in the Δ*gshF* and ΔPDE Δ*gshF* strains, which cannot synthesize GSH. These strains were grown in LSM until mid-exponential phase, then supplemented with GSH for ∼0.7 doubling times, and quantified for cytoplasmic GSH levels. In these assays, we found that ΔPDE Δ*gshF* accumulated about one third of GSH levels compared to Δ*gshF*, suggesting a GSH uptake defect at high c-di-AMP levels (**Fig. 4A**). We further verified this phenotype in ^3^H-GSH uptake assays for the Δ*gshF* and ΔPDE Δ*gshF* cultures (**Fig. 4B**).

**Figure 4:**
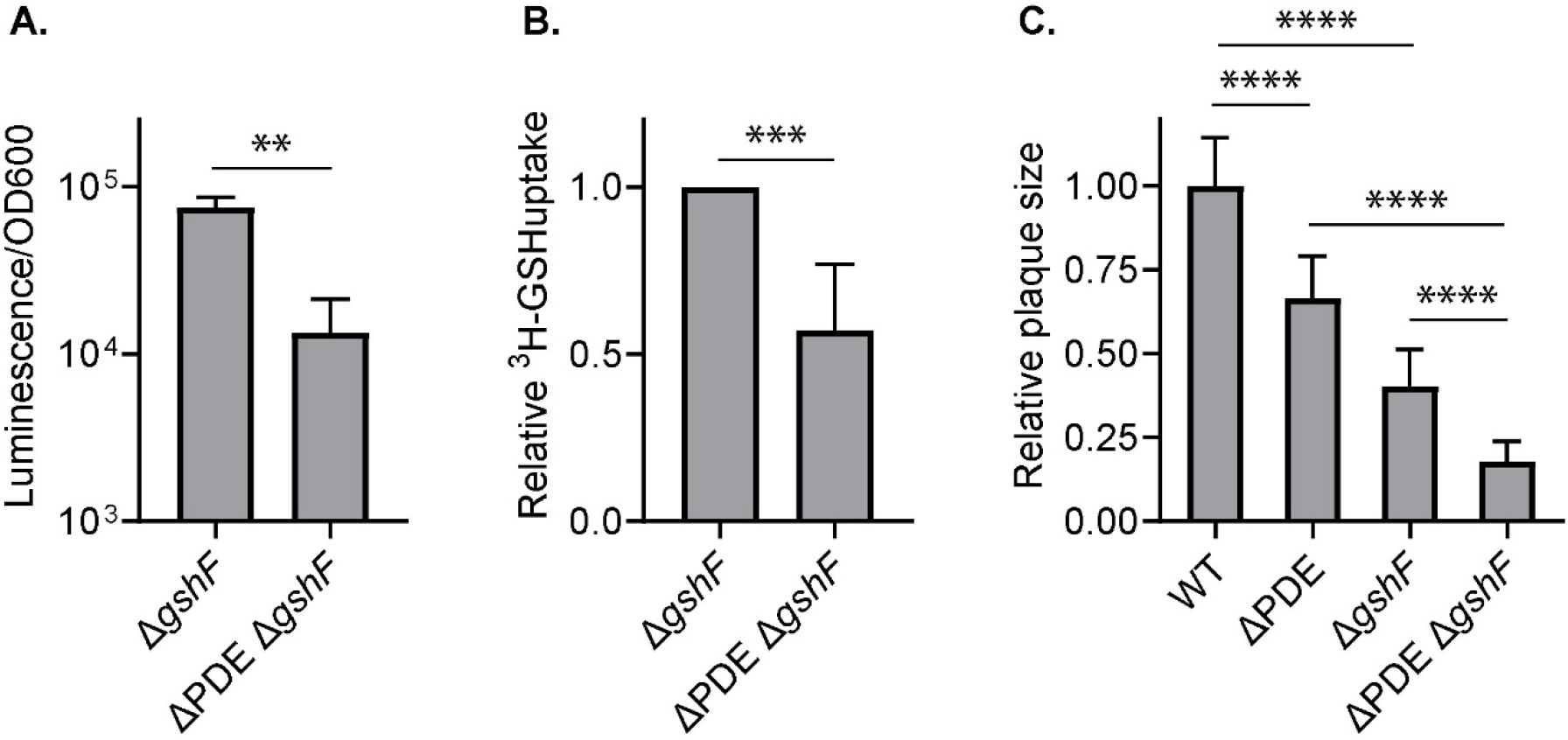
C-di-AMP accumulation inhibits GSH uptake in broth culture and ex-vivo infection. **A**. The Δ*gshF* and ΔPDE Δ*gshF* strains were grown to mid-log phase in LSM and supplemented with 10mM GSH for one hour (approximately 0.7 doublings in LSM). Cultures were thoroughly washed and quantified for intracellular GSH, reported as luminescence normalized to OD600. **B**. ^3^H-GSH uptake assay. Cultures were grown to mid-log, then supplemented with ^3^H-GSH for 30 minutes. Uptake activity was assessed by radioactivity normalized to OD600. **C**. Plaque formation at 4 days post infection of L2 fibroblasts. Plaque sizes were quantified by ImageJ. In each experiment, the average plaque size by each strain was normalized to the average WT plaque size, set at 1. Error bars represent standard deviations. Statistical analyses were performed by Student’s t-test in A-B and one-way ANOVA in C, with multiple comparisons for the indicated pairs: ^**^, P < 0.01; ^***^, P < 0.001; ^****^, P < 0.0001

The intracellular lifecycle of *L. monocytogenes*, including cytosolic replication and cell-to-cell spread, can be assessed by a plaque formation assay upon infection of murine fibroblasts (L2 cells) (27). To evaluate GSH uptake during infection, we performed an L2 plaque formation assay. Consistent with a previous study (5), we found the Δ*gshF* mutant to form significantly smaller plaques than the WT strain (**Fig. 4C**). Compared to Δ*gshF*, the ΔPDE Δ*gshF* strain was much further diminished for plaque formation, indicating that c-di-AMP accumulation impairs GSH uptake during infection (**Fig. 4C**).

### C-di-AMP also depletes cytoplasmic GSH by inhibiting synthesis or promoting catabolism

The ΔPDE mutant was deficient in intracellular GSH during growth in the absence of GSH supplementation, suggesting a defect in GSH synthesis (**Fig. 3A**). We first quantified glycine, cysteine, and glutamate, which are substrates for GSH synthesis by GshF (28). By LC-MS, we found that the ΔPDE strain was moderately reduced in glutamate, but glycine and cysteine levels were comparable to WT (**Fig. S3A**). Given that cysteine is the limiting substrate for GshF activity in bacterial cells (29), it is unlikely that the ΔPDE strain is lacking in substrates for GSH synthesis.

The gene expression levels of *gshF* were comparable in the WT and ΔPDE strains, with and without exogenous GSH in the culture medium (**Fig. S3B**). Therefore, we next examined post-transcriptional regulation of GshF by replacing the native *gshF* gene with a constitutively expressed allele (P_spac_-*gshF*). To evaluate GSH synthesis, we grew cultures in the absence or presence of TCEP, which activates GSH synthesis without adding GSH to the culture medium (8). Even upon constitutive *gshF* expression, the ΔPDE mutant was still significantly depleted for GSH, both under the basal condition and TCEP activation (**Fig. 5A**). In L2 plaque formation assay, *gshF* over-expression only moderately increased the ΔPDE plaque sizes, suggesting that GshF activity might be impaired (**Fig. 5B**). However, we noticed that the ΔPDE Δ*gshF* strain formed significantly smaller plaques than the ΔPDE strain, indicating that GSH synthesis is not completely inhibited at high c-di-AMP levels during infection (**Fig. 4C**). Therefore, c-di-AMP accumulation might also promote GSH catabolism.

**Figure 5:**
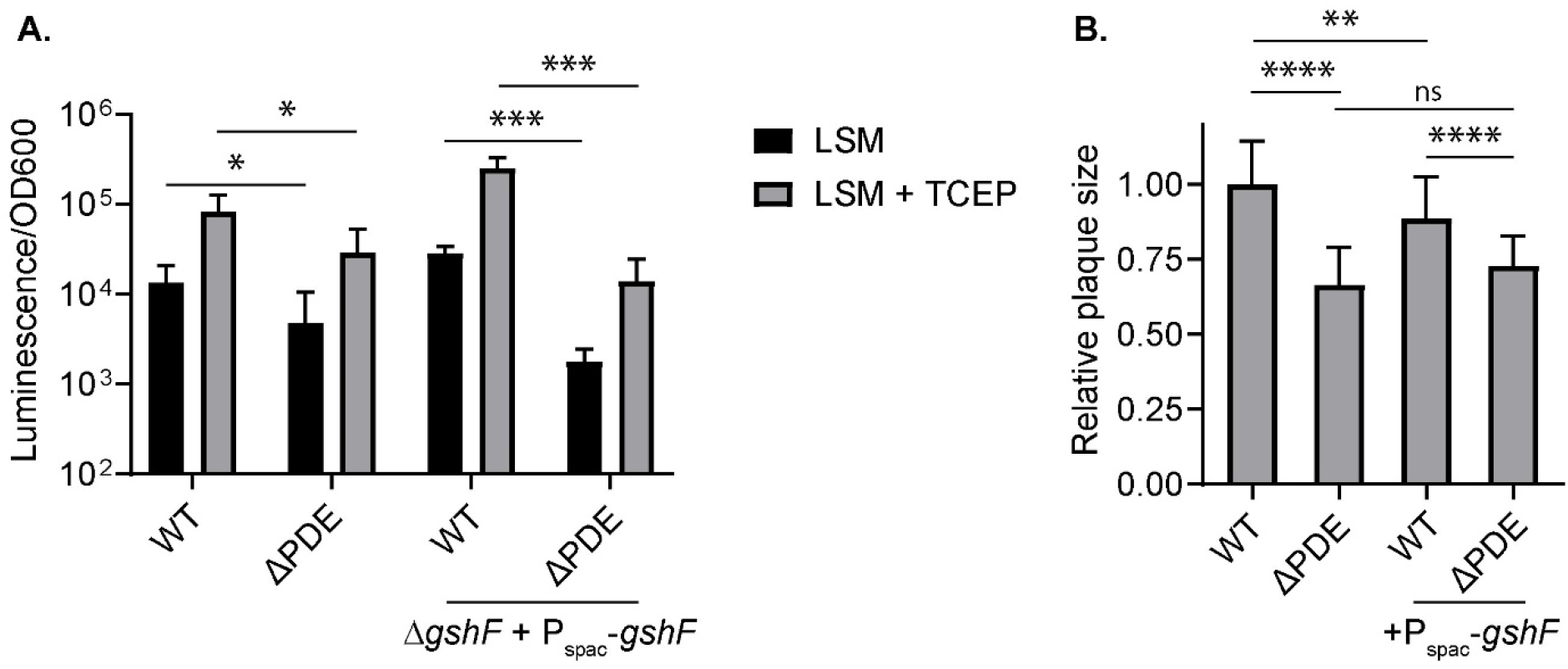
Activation of GSH synthesis activity does not restore GSH levels in the ΔPDE strain. **A**. Intracellular GSH levels. Cultures were left untreated or treated with 2mM TCEP for 2 hours. GSH levels are reported as luminescence normalized to OD600 of each culture. **B**. Plaque formation at 4 days post infection of L2 fibroblasts. Plaque sizes were quantified by ImageJ. In each experiment, the average plaque size by each strain was normalized to the average WT plaque size, set at 1. Error bars show standard deviations. Statistical analyses were performed by two-way ANOVA in A and one-way ANOVA in B, with multiple comparisons for the indicated pairs: ns, non-significant; ^*^, P < 0.05; ^**^, P < 0.01; ^***^, P < 0.001; ^****^, P < 0.0001

## DISCUSSION

C-di-AMP homeostasis is critical for bacterial growth and pathogenesis. The attenuated virulence conferred by c-di-AMP accumulation is widely observed in many pathogens, including *Mycobacterium tuberculosis*, which does not require c-di-AMP for growth (30, 31), and *Borrelia burgdorferi*, which produces a very low level of c-di-AMP (32, 33). In the c-di-AMP phosphodiesterase (Δ*dhhP*) mutant of *B. burgdorferi*, the transcription factor BosR is inhibited, causing reduced expression of the major virulence factor OspC (32). However, the *B. burgdorferi* Δ*dhhP* mutant also has a significant growth defect, and DhhP might degrade other nucleotides in addition to c-di-AMP (34, 35). For *Bacillus anthracis*, the ΔPDE mutant also grows poorly and is diminished for anthrax toxin production, a defect that is partially attributed to the upregulation of the transcriptional repressor AbrB (36). The molecular mechanisms by which c-di-AMP inhibits virulence factor expression in these bacteria are unclear, and the mechanisms by which c-di-AMP impairs the virulence of other pathogens are even less understood.

The *L. monocytogenes* c-di-AMP ΔPDE mutant does not have a significant growth defect in rich media, but is attenuated for virulence, suggesting defects that are pertinent to replication and adaptation in the host. Our findings here reveal GSH metabolism as another pathway that is regulated by c-di-AMP with implications for *L. monocytogenes* pathogenesis. First, we found that c-di-AMP accumulation inhibits GSH uptake. High affinity GSH transport in *L. monocytogenes* was recently shown to require the Ctp complex and the ATPases OppDF, the latter of which are components of the Opp complex that transports oligopeptides. Based on the demonstrated function of c-di-AMP in inhibiting potassium, Mg^2+^, and carnitine transporters (10), it is conceivable that c-di-AMP might also inhibit the OppDF and Ctp ATPase function particularly. As for how c-di-AMP accumulation depletes cytoplasmic GSH, we found that c-di-AMP does not inhibit *gshF* gene expression. Although our ex vivo infection data suggests that GshF is operational in the ΔPDE strain during infection, its activity might be reduced. Finally, c-di-AMP might also activate GSH catabolism. Future investigations are necessary to distinguish these mechanisms.

Because GSH is an allosteric activator of the master virulence factor PrfA, the first major implication of GSH depletion is a reduced virulence program expression. In support for this, our data shows that virulence gene expression at high c-di-AMP levels can be restored by a constitutive PrfA^*^ variant, or a combination of increased GSH synthesis and GSH supplementation. However, the PrfA^*^ variant does not fully rescue ΔPDE virulence to the WT level, indicating that c-di-AMP accumulation confers additional defects during infection. We previously reported that the *L. monocytogenes* ΔPDE strain is sensitive to cell wall-targeting antimicrobials, including lysozyme that is abundant in mammalian cell cytosol (37). Given the requirement of cell wall homeostasis for successful infection by *L. monocytogenes* (38–40), it is conceivable that cell wall defects also contribute to ΔPDE virulence attenuation. Another well-established function of c-di-AMP in *L. monocytogenes* is the inhibition of the carnitine transporter OpuC and multiple potassium uptake channels (12, 13). However, mutants abolished for carnitine transport do not exhibit a systemic infection defect like the ΔPDE strain (41), and the cell cytosol is highly enriched in potassium (42). Therefore, reduced osmolyte uptake may not significantly impact *L. monocytogenes* replication in the cytosol. Among other c-di-AMP-binding proteins that have been studied in *L. monocytogenes*, PstA has a minor role in infection (14); and the ΔPDE mutant does not have a growth defect that would be associated with a significantly reduced pyruvate carboxylase function (11, 43). The roles of other c-di-AMP molecular targets in *L. monocytogenes* pathogenesis should be investigated in future studies.

In addition to recognizing GSH as a virulence signal, *L. monocytogenes* also metabolizes GSH as a source of essential cysteine and uses GSH as antioxidant in broth cultures (26, 44). If these functions are relevant in the mammalian host, they can also explain why the ΔPDE *prfA*^*^ strain is not fully restored for virulence. Furthermore, a reduced function of the GSH importers Ctp and OppDF at high c-di-AMP levels can also deprive *L. monocytogenes* of nutrient oligopeptides, which are also imported through these uptake channels (45, 46).

It is curious that the PrfA^*^ (G145S) variant, which is active independent of GSH binding, does not confer constitutive *actA* expression in ΔPDE in the absence of GSH supplementation. Outside of the allosteric activation site that binds GSH, PrfA has four cysteine residues that can be glutathionylated (5). Although this modification modestly contributes to PrfA activation in the WT strain, it might have a more significant role at high c-di-AMP levels. Furthermore, the regulatory mechanisms of PrfA activity are complex, and PrfA can be inhibited by several factors such as metals or oligopeptides that do not contain cysteine (4). It is conceivable that the ΔPDE strain accumulates those PrfA inhibitors that can be identified in future metabolomics studies.

Altogether, our studies present an exciting avenue to investigate how c-di-AMP disrupts GSH metabolism in *L. monocytogenes*. Given the importance of GSH as a virulence signal, nutrient, and antioxidant in different bacteria (47), it will be interesting to examine whether c-di-AMP regulates GSH metabolism in other species, and the consequences of such regulation on bacterial growth and infection.

## MATERIALS AND METHODS

### Strains and culture conditions

*L. monocytogenes* strains are listed in **Table S2**. Improved *Listeria* Synthetic Medium (LSM) was prepared based on published recipe (20). Over-expression of genes was accomplished using the integrative plasmid pPL2 (48). Gene allelic exchange was performed using plasmid pKSV7. For all experiments related to broth cultures, *L. monocytogenes* was grown at 37°C with shaking and without antibiotics. For glycerol stocks and maintenance, strains carrying derivatives of the integrative plasmid pPL2 were grown in Brain Heart Infusion broth with 200μg/mL streptomycin and 10μg/mL chloramphenicol. For mouse and L2 cell infection, *L. monocytogenes* was grown static in BHI at 30°C.

### RNAseq and data analysis

*L. monocytogenes* cultures were grown to OD_600_ of ∼ 0.5, and harvested by mixing 1:1 with cold methanol and centrifugation. RNA was extracted using acidified phenol:chloroform at pH 5.2 as previously described (49). Contaminated DNA was removed from extracted RNA using Turbo DNase (Thermo Scientific). RNA sequencing and read mapping was performed by MiGS Sequencing Center (Pittsburgh, PA). Reads were mapped to the *L. monocytogenes* 10403S genome (NCBI:txid393133). Differentially expressed gene analysis was performed by EdgeR (50).

### RT-qPCR for *L. monocytogenes* genes in broth

For quantification of PrfA activity upon activation by GSH, *L. monocytogenes* cultures were grown in LSM or LSM + 10mM GSH to OD_600_ ∼0.5. For PrfA activation by TCEP, *L. monocytogenes* cultures were grown in LSM to OD ∼0.5, then split into two aliquots of identical volumes. One culture aliquot was left untreated and the other was supplemented with 2mM TCEP for 2 hours, and both were shaken at 37°C. RNA was extracted, treated with Turbo DNase and converted to cDNA using iScript cDNA synthesis kit (BioRad). Gene expression was quantified using iTaq Universal SYBR Green (BioRad) with primers specific to each target, with *rplD* as the control housekeeping gene. Primers are listed in **Table S3**.

### Quantification of GSH and GSSG

*L. monocytogenes* cultures were grown to an OD_600_ of ∼0.5 and harvested by centrifugation. Cell pellets were washed three times in phosphate-buffered saline, and lysed by sonication. Lysates were centrifuged to remove cell debris. GSH and GSSG were quantified using the luminescence-based GSH-Glo assay kit (Promega). GSH standards of 0.5 – 20 µM were quantified to determine the linear range of luminescence readings, and bacterial lysates were diluted appropriately such that luminescence values were within the linear range.

### L2 plaque assay

Plaque formation upon *L. monocytogenes* infection of L2 cells was quantified as previously described (27, 49). Briefly, 1.2x10^6^ cells were infected with *L. monocytogenes* at multiplicity of infection (MOI) of 0.25. At 1 hour post infection, cells were washed, and fresh cell medium in 0.7% agarose was supplemented with 10µg/mL gentamicin to kill extracellular *L. monocytogenes*. At 5 days post infection, cells were stained with 0.3% crystal violet to visualize plaques. Plaque sizes were analyzed using ImageJ software.

### P_actA_::RFP assay

A pPL2 plasmid expressing red fluorescent protein from the *actA* promoter was used to quantify transcriptional activity. Fluorescence assays were performed based on published methods (8). Briefly, *L. monocytogenes* strains carrying pPL2-P_actA_::RFP were grown in overnight at 37°C in LSM. Cultures were diluted 1:10 into fresh LSM, LSM + 10mM GSH, or LSM + 2mM TCEP and grown to OD_600_ of 1.5-2. Cultures were harvested by centrifugation and resuspended in phosphate-buffered saline to achieve a 5X concentration. Fluorescence was quantified in black-bottom 96-well plates (Greiner Bio) for fluorescence measurement (excitation 540nm, emission 650nm) (BioTek Synergy H1 Microplate Reader). Fluorescence values were normalized to OD_600_.

### Mouse infection

All techniques were reviewed and approved by the University of Wisconsin—Madison Institutional Animal Care and Use Committee under the protocol M005916-R01-A01. Five 6-week-old female C57BL/6 mice (Charles River NCI facility) per bacterial strain were infected via tail vein injection with 0.2 cc of PBS with 1x10^5^ of logarithmically growing bacteria. At 48hpi, spleens and livers were harvested, and homogenized in organ lysis buffer. Organ homogenates were diluted, as necessary using PBS, spiral plated on LB agar, and CFU counts were obtained using the Q count. Bacterial CFUs per organ were calculated using known dilution factors.

### Quantification of amino acids by LC-MS

*L. monocytogenes* strains were grown overnight in 3mL LSM cultures at 37°C, shaking. Overnight cultures were diluted 1:100 in LSM and grown with aeration to mid logarithmic phase (0.4-0.6 OD_600_) at 37°C, shaking at 200 rpm. Bacterial metabolite extraction and quantification were performed as previously described (39). Briefly, bacterial cells were filtered and resuspended in extraction solvent (HPLC-grade acetonitrile, methanol, and water at a 2:2:1 ratio), and immediately frozen on in dry ice. Samples were analyzed on a Dionex UHPLC system coupled with a hybrid quadrupole–high-resolution mass spectrometer (Q Exactive Orbitrap; Thermo Scientific), using an Acquity UPLC BEH C18 column (Waters). Solvent A was composed of 97% water, 3% methanol, 10 mM tributylamine, pH 8.1-8.2. Solvent B was 100% methanol. Mobile phase was run on a gradient from 5 – 95% solvent B over 25 minutes. MS data in mzXML format was used for metabolite identification with the MAVEN software (51)

## ACKNOWLEDGEMENTS

This work was supported by R35 GM147519 (TNH) and R01 AI137070 (JDS).

